# Development and verification of SSR markers from drought stress-responsive miRNAs in common wild rice

**DOI:** 10.1101/2021.10.08.463621

**Authors:** Yong Chen, Yuan-wei Fan, Wan-ling Yang, Gu-mu Ding, Min-min Zhao, Yan-hong Chen, Jian-kun Xie, Fan-tao Zhang

## Abstract

**PREMISE:** Dongxiang wild rice *(Oryza rufipogon* Griff., DXWR) is the northernmost common wild rice found in the world, which possesses abundant elite genetic resources. We developed a set of drought stress-responsive microRNA (miRNA)-based single sequence repeat (SSR) markers for DXWR, which will help breed drought stress-resistant rice varieties.

**METHODS AND RESULTS:** Ninety-nine SSR markers were developed from the drought stress-responsive miRNAs of DXWR. The SSR loci were distributed in all 12 rice chromosomes and most were in chromosomes 2 and 6, with di- and trinucleotides being the most abundant repeat motifs. Nine out of ten synthesized SSR markers were displayed high levels of genetic diversity in the genomes of DXWR and 41 modern rice varieties worldwide. The number of alleles per locus ranged from 2 to 6, and the observed and expected heterozygosity ranged from 0.000 to 0.024 and 0.461 to 0.738, respectively.

**CONCLUSIONS:** These SSR markers developed from drought stress-responsive miRNAs in DXWR could be additional tools for elite genes mapping and useful for drought stress-resistant rice breeding.

---

Rice (*Oryza sativa* L.) is the staple food for half of the world’s population and it has shaped billions of people’s diets, economies, and cultures (Zhu et al., 2017). As the world population grows constantly, so does the demand for rice (Feng et al., 2021). However, securing rice production has become a great challenge because it is usually affected by various biotic and abiotic stresses (Liu et al., 2019). It has been estimated that rice production may decline by 3.8% in the Asian region in 21 century due to various stresses (Matthews et al., 1997). Among them, drought stress is one of the biggest factors seriously affecting rice growth and yield, because rice is a semi-aquatic plant (Berahim et al., 2021). Rice plants need more water during their whole life than other cereal crops, such as maize and wheat. Normally, early drought stress results in delayed transplantation, affecting seed germination and seedling growth, finally ending with harvest failure. During the rice reproductive stage, drought stress causes grain sterility and poor grain filling. Therefore, identification of drought stress-resistant gene resources and the subsequent development of drought stress-resistant rice varieties are of great importance for rice production.

Common wild rice (*Oryza rufipogon* Griff.) is the ancestor of modern rice varieties and possesses many valuable gene resources that have been lost during the process of domestication (Shishido et al., 2019). Dongxiang wild rice (DXWR) is the northernmost (28°14’ N) common wild rice in the world and has strong drought stress resistance (Zhang et al., 2016). Therefore, DXWR is an elite germplasm resource for the development and breeding of new drought stress-resistant rice varieties. MicroRNAs (miRNAs) are a class of non-coding RNAs (approximately 21 bp long) that can regulate gene expression at the post-transcriptional level in plants (Zhang et al., 2014). A considerable number of studies have shown that miRNAs play vital roles in drought stress resistance in many plant species, including sugarcane (Gentile et al., 2015), soybean (Li et al., 2016), wheat (Akdogan et al., 2016), rice (Nadarajah and Kumar, 2019), *Arabidopsis* (Niu et al., 2019), and tomato (Zhou et al., 2020). In our previous study, we found that miRNAs also play crucial roles in response to drought stress in DXWR, and 33 conserved miRNAs and 67 novel miRNAs were detected to be significantly affected by drought stress (Zhang et al., 2016; Zhang et al., 2018).

Developing drought stress-resistant plants with the aid of gene marker-assisted breeding has provided great benefits for conventional breeding (Luo et al., 2019). Simple sequence repeats (SSRs), also called microsatellites, are DNA fragments with 1-6 nucleotide motifs repeated a variable number of times. SSRs are ideal genetic markers because they have many advantages, such as genetic co-dominance, abundance, rich polymorphism, high reliability, and dense genome-wide coverage (Li et al., 2019). Consequently, SSR markers are considered to be powerful tools for evaluation of genetic diversity, germplasm resources identification, and genetic mapping. To date, some studies have developed SSR markers based on miRNAs in plants. For example, Mondal and Ganie (2014) developed a set of SSR markers from 130 members of salt stress-responsive miRNAs in rice. Sihag et al. (2021) developed 70 miRNA-based SSR markers using a set of 20 terminal heat stress-tolerant and heat stress-susceptible wheat genotypes. We recently developed the IncRNA-derived SSR markers. However, to our knowledge, there are few studies on the development of miRNA-based SSR markers in common wild rice, which has strong resistance to various abiotic stresses. Therefore, the present study was aimed at the development of a set of additional promising SSR markers from drought stress-responsive miRNAs for the elite genes mapping and molecular breeding programs of DXWR.

## METHODS AND RESULTS

Previously, we identified 100 drought stress-responsive miRNAs (33 conserved and 67 novel) in DXWR and obtained 99 corresponding pre-miRNA sequences through high-throughput sequencing approaches, because two conserved miRNAs (miR2871a-3p and miR2871a-5p) shared the same pre-miRNA sequences. In this study, the 99 pre-miRNA sequences were blasted with the Nipponbare MSU v7 reference genome (http://rice.uga.edu/) to obtain their flanking sequences, which included 500 bp upstream and 500 bp downstream regions of the pre-miRNA sequences. Finally, we obtained 99 sequences that contain 100 drought stress-responsive miRNAs and their corresponding flanking sequences. Then, the 99 sequences were screened by the SSRHunter software to find SSR loci with a 4 bp motif maximum and three repeat minimum (Li and Wan, 2005). A total of 673 miRNA-based SSRs were obtained in all sequences. Then, primer pairs of each of the SSR-containing sequences were designed using the program Primer3 Input (version 0.4.0) (https://bioinfo.ut.ee/primer3-0.4.0/). The parameters were set using the following criteria: (1) optimum primer length of 18-23 bp, (2) GC content of 40%-60%, (3) annealing temperature between 50°C and 60°C (55°C optimum), and (4) expected amplicon size of 100-300 bp; the rest of the parameters were set as defaults.

Finally, we developed 99 primer pairs for the 100 drought stress-responsive miRNAs of DXWR. These markers were distributed in all of the 12 chromosomes and were most abundant in chromosomes 2 and 6 (11), and chromosomes 1, 4 and 8 (10) (Table 1). Among them, dinucleotide (56, 56.6%) and trinucleotide (36, 36.4%) repeat motifs were the most abundant (Table 1). Meanwhile, the AT was the most abundant (21.4%) type of dinucleotide repeats, followed by AG (14.3%) and GA (12.5%) types (Table 1).

**TABLE 1.**
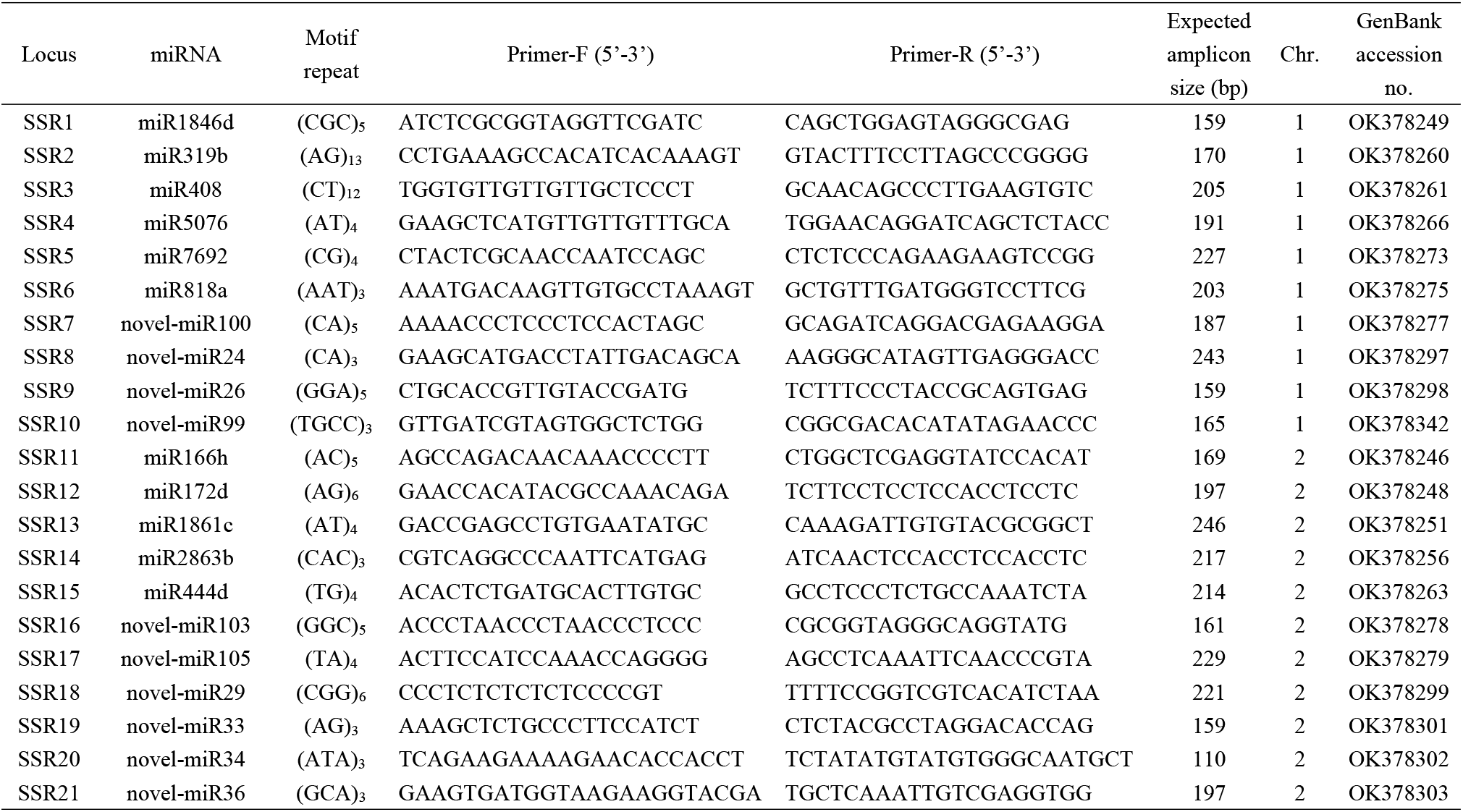
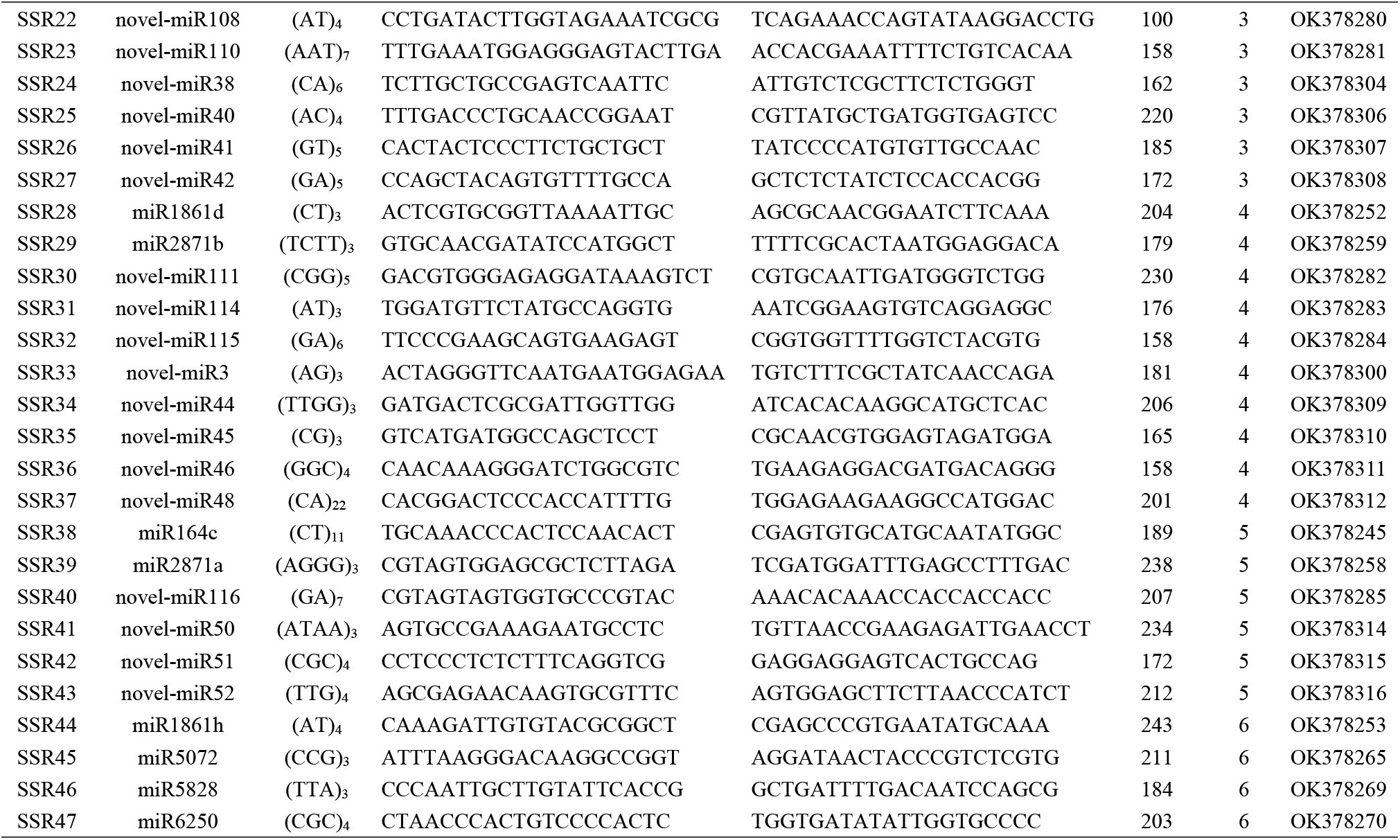
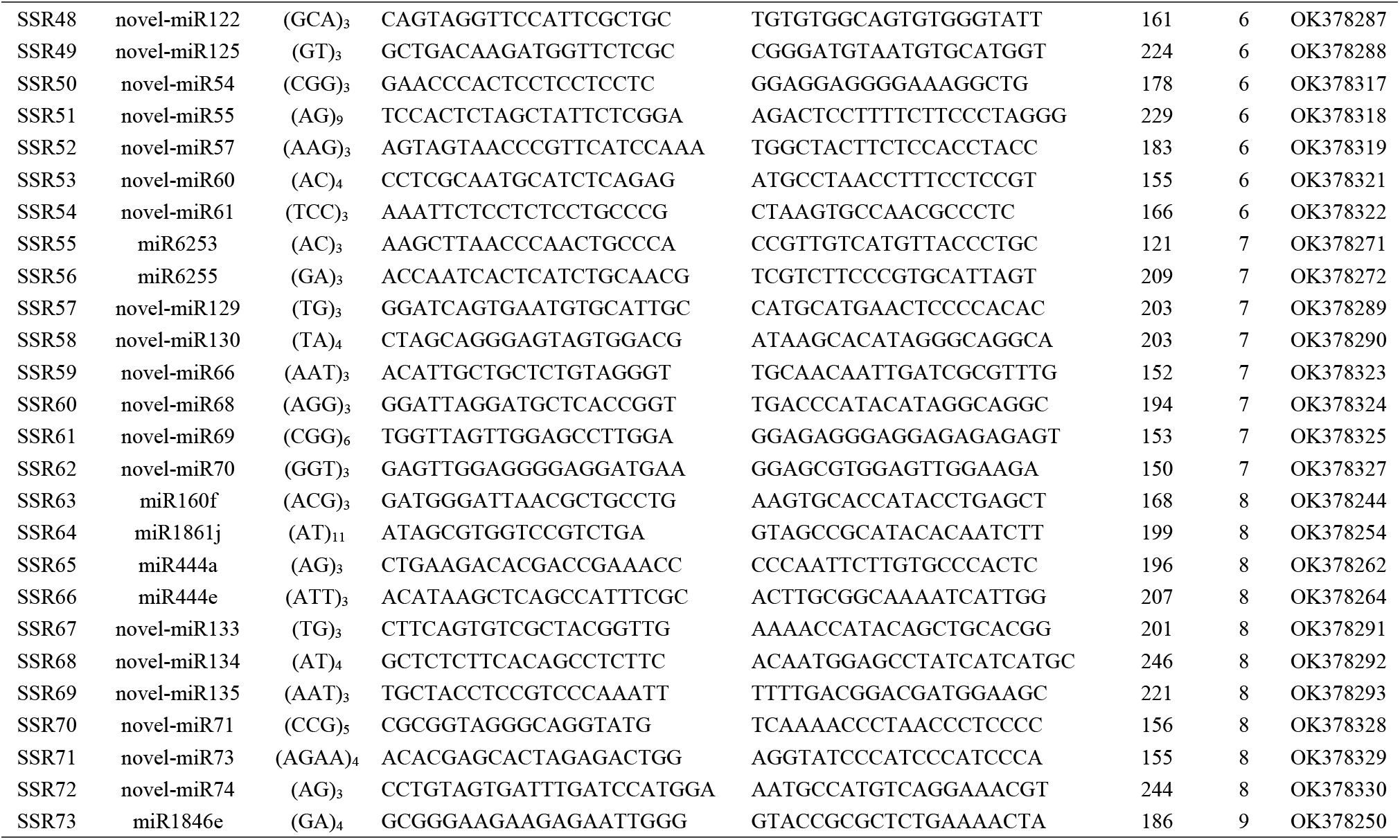
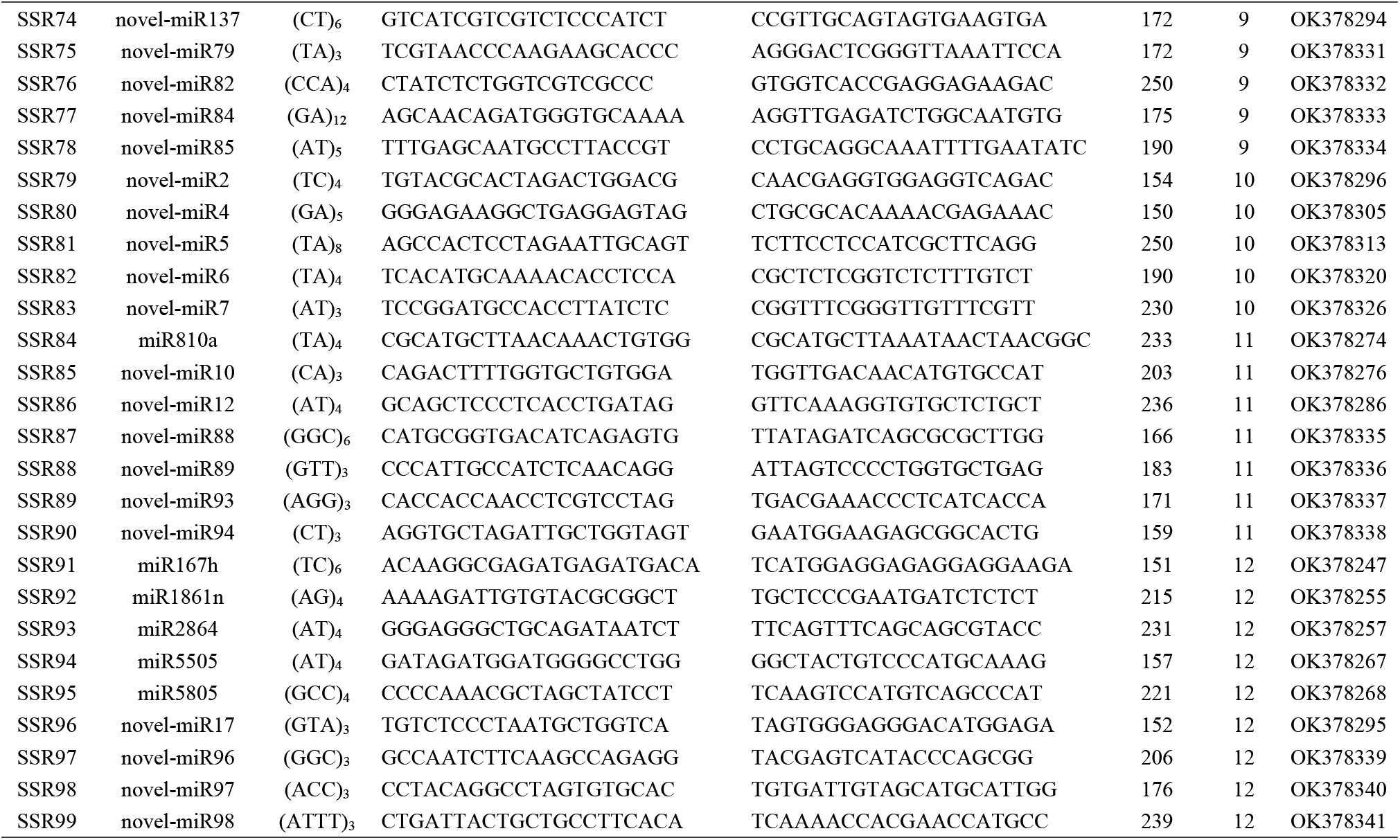
Characteristics of SSR markers derived from the drought stress-responsive miRNAs of DXWR.

To investigate whether these miRNA-based SSR markers developed for DXWR could be applied to the modern rice varieties, we synthesized 10 pairs of primers and then amplified fragments using the genomic DNA of DXWR and 41 modern rice varieties worldwide. Among these rice varieties, 18 belonged to *indica*, and the remaining 23 belonged to *japonica* (Appendix 1). Fresh leaf tissues were used for total genomic DNA extraction using a Rapid Plant Genomic DNA Isolation Kit (Sangon Biotech, Shanghai, China). PCR amplifications were carried out in a 10 μL reaction system containing 1 μL (200 ng/μL) genomic DNA, 1 μL (0.01 nmol/μL) primers, 3 μL ddH2O, and 5 μL 2 × Fast Taq Premix (Tolo Biotech Co., Ltd, China). The PCR reaction conditions were as follows: initial denaturation at 95°C for 5 min; 35 cycles of denaturation at 95°C for 30 s, annealing at 56°C for 30 s, extension at 72°C for 30 s, and 5 min at 72°C for the final extension. The PCR products were separated on 6% denaturing polyacrylamide gels in 0.5 × TBE buffer and the bands were visualized via silver nitrate staining. All of the 10 primer pairs were amplified successfully for DXWR and the modern rice varieties. The electrophoresis results suggested that nine out of the 10 primer pairs were polymorphic among DXWR and the modern rice accessions. The nine polymorphic SSR markers were analyzed using GenAlEx 6.5 software for the number of alleles per locus, observed heterozygosity, and expected heterozygosity (Peakall and Smouse, 2012). Software Arlequin ver 3.5 (Excoffier and Heidi, 2010) was used to determine Hardy-Weinberg equilibrium (HWE). Among the nine polymorphic loci, the number of alleles per locus ranged from two to six with a mean of 4.444 (Table 2). The levels of observed and expected heterozygosity were 0.000-0.024 and 0.461-0.738, with averages of 0.003 and 0.625, respectively (Table 2). Among the nine polymorphic primer pairs, seven pairs were able to detect levels of expected heterozygosity above 0.5, indicating a high level of polymorphism among these rice accessions (Table 2).

**TABLE 2.** Genetic properties of the 9 polymorphic SSR markers among DXWR and 41 modern rice varieties.

| Locus | miRNA | $A$ | $H_o$ | $H_e^a$ |
| --- | --- | --- | --- | --- |
| SSR2 | miR319b | 2.000 | 0.000 | 0.672** |
| SSR3 | miR408 | 5.000 | 0.000 | 0.461** |
| SSR23 | novel-miR110 | 5.000 | 0.000 | 0.738** |
| SSR37 | novel-miR48 | 6.000 | 0.024 | 0.683** |
| SSR38 | miR164c | 4.000 | 0.000 | 0.673** |
| SSR51 | novel-miR55 | 4.000 | 0.000 | 0.622** |
| SSR64 | miR1861j | 5.000 | 0.000 | 0.498** |
| SSR77 | novel-miR84 | 6.000 | 0.000 | 0.622** |
| SSR81 | novel-miR5 | 3.000 | 0.000 | 0.656** |
| Mean | - | 4.444 | 0.003 | 0.625 |

*Note:* $A$ = number of alleles per locus; $H_o$ = observed heterozygosity; $H_e$ = expected heterozygosity.
<sup>a</sup> Deviations from Hardy-Weinberg equilibrium: \*\* $P < 0.01$ .

## CONCLUSIONS

In this study, we developed a set of SSR markers for the drought stress-responsive miRNAs in DXWR and verified the validity and applicability of these valuable markers in rice varieties grown worldwide. This is the first report on the development and verification of miRNA-based markers for abiotic stress resistance in DXWR, which lays the foundation for future basic and applied research to make full use of elite miRNA genes from this valuable wild rice germplasm resources.

## ACKNOWLEDGEMENTS

This research was partially supported by the National Natural Science Foundation of China (31960370, 32070374, 31960085), the Natural Science Foundation of Jiangxi Province, China (20202ACB205002), the Foundation of Jiangxi Provincial Key Lab of Protection and Utilization of Subtropical Plant Resources (YRD201913), and the Postgraduate Innovation Fund of Jiangxi Normal University (YC2019-B043).

## AUTHOR CONTRIBUTIONS

F.-T.Z., and J.-K.X. conceived and designed the experiments. W.-L.Y., and G.-M.D. collected the samples. Y.C., and W.-L.Y., designed the primers. Y.C., and Y.-H.C. performed the molecular laboratory work. Y.C. analyzed the data. Y.C., Y.-W.F., and F.-T.Z. drafted the manuscript.

## DATA AVAILABILITY STATEMENT

All of the sequences of microsatellite loci had been deposited into the National Center for Biotechnology Information (NCBI)’s GenBank database with the accession number of OK378244 – OK378342.

## APPENDIX 1. Worldwide rice cultivars used for verification of the developed SSR markers in this study

|  | Name | Origin | Subspecies |
| --- | --- | --- | --- |
| 1 | Xieqingzao B | China | *indica* |
| 2 | Ai-Chiao-Hong | China | *indica* |
| 3 | Ming Hui | China | *indica* |
| 4 | ZHE 733 | China | *indica* |
| 5 | Zhenshan 2 | China | *indica* |
| 6 | Aijiaonante | China | *indica* |
| 7 | Hon Chim | Hongkong, China | *indica* |
| 8 | Binulawan | Philippines | *indica* |
| 9 | IR64 | Philippines | *indica* |
| 10 | Byakkoku Y 5006 Seln | Australia | *indica* |
| 11 | Champa Tong 54 | Thailand | *indica* |
| 12 | Chau | Vietnam | *indica* |
| 13 | Darmali | Nepal | *indica* |
| 14 | Oryzica Llanos 5 | Colombia | *indica* |
| 15 | Criollo La Fria | Venezuela | *indica* |
| 16 | Radin Ebos 33 | Malaysia | *indica* |
| 17 | Tchibanga | Gabon | *indica* |
| 18 | SL 22-613 | Sierra Leone | *indica* |
| 19 | Zhonghua 11 | China | *japonica* |
| 20 | Shangyu 394 | China | *japonica* |
| 21 | Sung Liao 2 | China | *japonica* |
| 22 | Caawa/Fortuna 6-103-15 | Taiwan, China | *japonica* |
| 23 | Aichi Asahi | Japan | *japonica* |
| 24 | Jouiku 393G | Japan | *japonica* |
| 25 | Kibi | Japan | *japonica* |
| 26 | Nipponbare | Japan | *japonica* |
| 27 | Hiderisirazu | Japan | *japonica* |
| 28 | Hatsunishiki | Japan | *japonica* |
| 29 | Edomen Scented | Japan | *japonica* |
| 30 | Rikuto Norin 21 | Japan | *japonica* |
| 31 | Shirogane | Japan | *japonica* |
| 32 | Kiuki No. 46 | Japan | *japonica* |
| 33 | Arias | Indonesia | *japonica* |
| 34 | Geumobyeo | Korea, South | *japonica* |
| 35 | Lemont | United States | *japonica* |
| 36 | Moroberekan | Guinea | *japonica* |
| 37 | Ostiglia | Italy | *japonica* |
| 38 | Bombon | Spain | *japonica* |
| 39 | Coppocina | Bulgaria | *japonica* |
| 40 | WC 4443 | Bolivia | *japonica* |
| 41 | Tropical Rice | Ecuador | *japonica* |

